# Dipolar extracellular potentials generated by axonal projections

**DOI:** 10.1101/109918

**Authors:** Thomas McColgan, Ji Liu, Paula T Kuokkanen, Catherine E Carr, Hermann Wagner, Richard Kempter

## Abstract

Extracellular field potentials (EFPs) are an important source of information in neuroscience, but their physiological basis is in many cases still a matter of debate. Axonal sources are typically discounted in modeling and data analysis because their contributions are assumed to be negligible. Here, we show experimentally and theoretically that contributions of axons to EFPs can be significant. Modeling action potentials propagating along axons, we showed that EFPs were prominent in the presence of a terminal zone where axons branch and terminate in close succession, as found in many brain regions. Our models predicted a dipolar far field and a polarity reversal at the center of the terminal zone. We confirmed these predictions using EFPs from the barn owl auditory brainstem where we recorded in nucleus laminaris using a multielectrode array. These results demonstrate that axonal terminal zones produce EFPs with considerable amplitude and spatial reach.

## Introduction

Extracellular field potentials (EFPs) are at the heart of many experimental approaches used to examine the inner workings of the brain. Types of EFPs include invasively recorded signals such as the electrocorticogram (ECoG), the local field potential (LFP), the current source density (CSD), and the multiunit activity (MUA) as well as the noninvasively recorded electroencephalogram (EEG) and the auditory brainstem response (ABR) (Nunez and Srinivasan, 2006; Brette and Destexhe, 2012). The origins of these signals, especially in cases in which the activity is not clearly attributable to a single cell, is a matter of debate (Buzsáki et al., 2012).

EFPs in the brain were long thought to be primarily of synaptic origin (Buzsáki et al., 2012). As a consequence, many modeling studies focussed on the extracellular fields induced by postsynaptic currents on the dendrites and the soma of a neuron (Holt and Koch, 1999; Lindén et al., 2010, 2011; Einevoll et al., 2013; Fernández-Ruiz et al., 2013). However, a number of recent data analyses and modeling efforts have revealed that active, non-synaptic membrane currents can play an important role in generating population-level EFPs (Ray and Maunsell, 2011; Belluscio et al., 2012; Schomburg et al., 2012; Reimann et al., 2013; Waldert et al., 2013; Anastassiou et al., 2015; Ness et al., 2016), including far reaching potentials detectable at the scalp (Teleńczuk et al., 2011, 2015). Currents from the axon are still thought to be so small as to be of minor importance for the EFP.

One of the reasons for the assumption that axonal currents contribute little to the EFP is that the far field of an action potential traveling along an idealized straight axon is quadrupolar, meaning that it decays faster with distance than synaptic sources, which are typically dipolar (Nunez and Srinivasan, 2006). Surprisingly, some experimental studies indicated that the EFP of axonal responses may also have a dipolar structure. For example, Blot and Barbour (2014) reported an EFP with a characteristic dipolar structure in the vicinity of cerebellar Purkinje cell axons; other studies (Swadlow and Gusev, 2000; Swadlow et al., 2002) showed that the axonal part of the EFP of thalamocortical afferents showed a polarity reversal associated with a dipolar source, and classical current source density studies found dipolar current distributions in axonal terminal zones in the cortex and lateral geniculate nucleus, and attributed these to axons because of conduction velocities (Mitzdorf and Singer, 1977, 1978; Mitzdorf, 1985). Here we show a strong dipolar, axonal field potential from the auditory brainstem of the barn owl.

The discrepancy between the quadrupolar structure of EFPs generated by idealized axons, and the experimentally observed dipolar structure raises the question of how axons are able to generate dipolar field potentials. We show how dipolar far fields in the EFP of axons can be explained by the axons’ anatomical structure. In particular, the branchings and terminations of axons in their terminal zone area deform the extracellular waveform (Plonsey, 1977; Gydikov and Trayanova, 1986; Gydikov et al., 1986), and can lead to a dipolar EFP structure. Axon bundles, sometimes called fascicles, exist throughout the peripheral and central nervous system and have such terminal zones (Nornes and Das, 1972; Goodman et al., 1984; Hentschel and van Ooyen, 1999; Kandel et al., 2000). The white matter of the mammalian brain can be viewed as an agglomeration of such fascicles (Schüz and Braitenberg, 2002). We therefore predict pronounced contributions of axon bundles to EFPs, which are neglected in current models.

In what follows, axonal contributions to the EFP are first investigated by a numerical model based on forward simulation (Holt and Koch, 1999; Gold et al., 2006). This first model includes a large-scale multi-compartment simulation (Rall, 1959; Jack et al., 1975; Abbott, 1992; Hines and Carnevale, 1997; Hines et al., 2009) of an axon population. We then outline the basic mechanisms by means of a second, analytically tractable, model of a generic axon bundle. Finally, we validate model predictions with data from multi-site in-vivo electrophysiological recordings from the barn owl auditory brain stem.

## Results

### Effects of axonal bifurcations and terminations on extracellular action potentials

To understand how the geometry of an axon affects the extracellular waveform associated with action potentials, we first numerically simulated single action potentials propagating along generic axons and calculated their contribution to the EFP (for details, see Materials and Methods). This was done for five scenarios: infinite axons, terminating axons, bifurcating axons, axons that bifurcate as well as terminate, and axon bundles (Figure 1). We began by simulating an infinitely long axon following a straight line path, neither bifurcating nor terminating (Figure 1A). The extracellular action potential has the characteristic triphasic shape. As the action potential travels from top to bottom, the waveform is translated in time with the conduction velocity, but is otherwise unchanged. The triphasic shape is also present in the spatial arrangement of currents at any given time, which is the reason for the quadrupolar EFP response traditionally assumed for axons.

**Figure 1:**
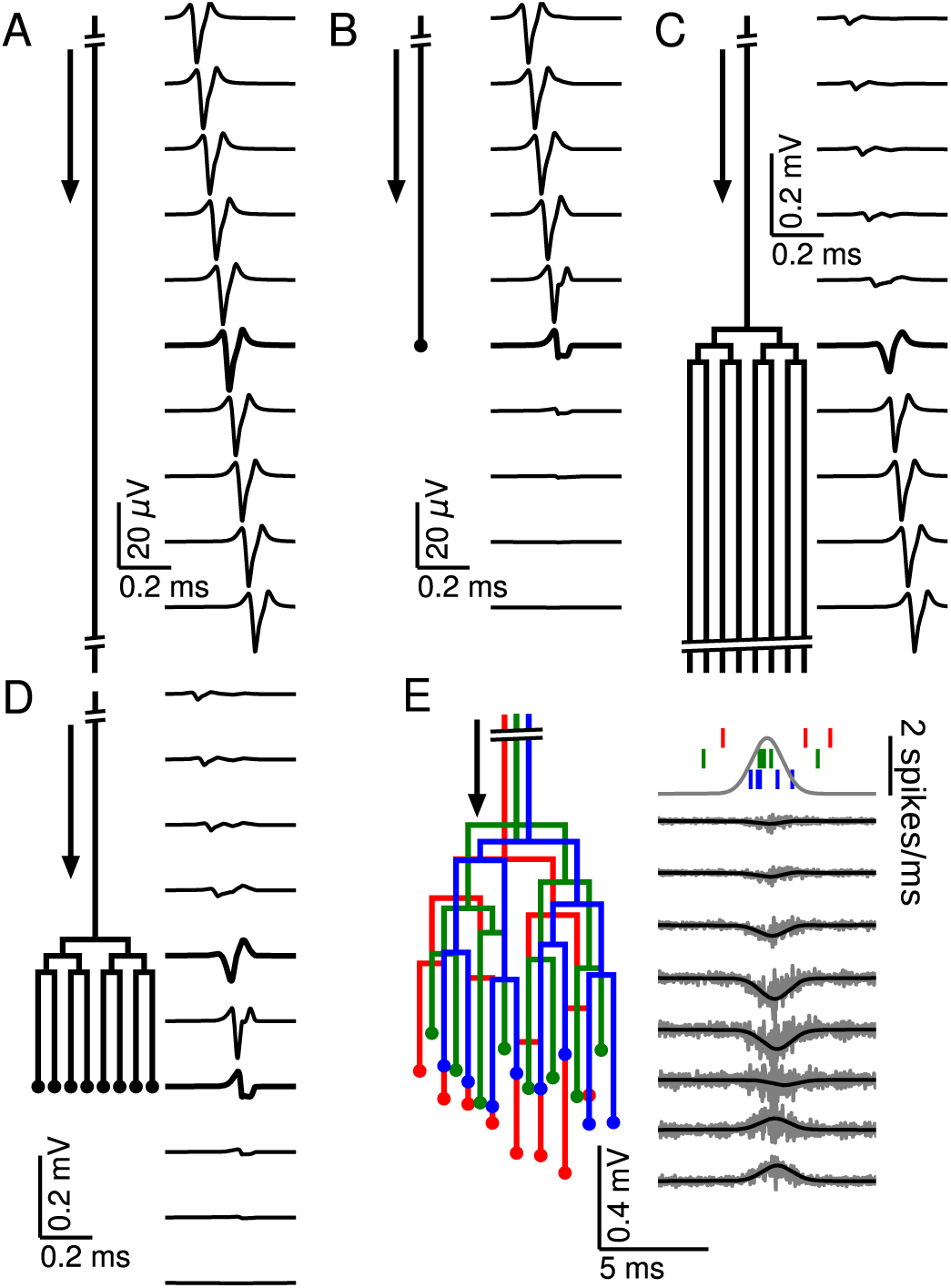
Relationship between axon morphology and extracellular potential. Multi-compartment simulations of action potentials traveling along axons with varying morphologies, as indicated by the diagram on the left-hand side of each subfigure. Action potential propagation direction indicated by arrow. Waveforms, shown on the right-hand side of each subfigure, were recorded at a horizontal distance of 150 μm from the axons. The vertical depth is indicated by the plot position, spaced by 200 μm. Horizontal distances between axons are for illustration only, all axons were simulated to lie on a straight line. (**A**) Action potential in an infinitely long, straight axon. (**B**) Terminating axon. Action potential waveform closest to the termination thickened for emphasis. (**C**) Branching axon. The axon branches multiple times within of 200 μm. Thicker waveform at the center of the bifurcation zone. (**D**) Combined bifurcations and terminations. Note the larger voltage scales in C and D, which correspond to the different number of fibers. (**E**) Response in a population of 100 randomized morphologies, three of which are shown schematically (colored). Activity consists of spontaneous background activity (100 spikes/s) superimposed with a brief Gaussian pulse of heightened spike rate (2000 spikes/s). Spike rate and example spike times for the three morphologies are shown at the top. Right: gray lines show activity of full population averaged over 40 trials, while the black lines show the low-pass (<1 kHz) component. Note that the time and voltage scales are different from A-D.

There are two ways of understanding the triphasic shape of the extracellular waveform. One way is by attribution of the peaks of the response to individual current types. The first, small positive peak corresponds to the capacitive current, the large and negative second peak to the sodium current, and the final positive peak to the potassium current (Gold et al., 2006). Another, more mathematical way of understanding the triphasic shape is specific to the nature of the axon. Due to Kirchhoff’s law and cable theory (see Materials and Methods), the local transmembrane current in a homogeneous axon is proportional to the second spatial derivative of the membrane potential along the direction of the axon. Because the action potential is roughly a traveling wave, the currents are also proportional to the second *temporal* derivative of the membrane potential. The three extrema of the EFP are thus related to the points of maximum curvature in the action potential waveform, namely the onset, the maximum, and the end of the spike.

Next we simulated the response of an axon that terminates (Figure 1B). Here the action potential approaching the recording location (top traces) has the same, triphasic EFP response as in the non-branching case. When the action potential reaches the termination point, its EFP gradually deforms into a biphasic response, with a positive peak preceding a negative peak. The mechanism for this deformation can be understood as follows: As the action potential approaches the recording location next to the termination, the majority of the transmembrane currents flow at points located before the termination, and they are almost identical to those in the non-terminating case; the first, capacitive peak is not affected. As already mentioned, the second and third peaks of the extracellular action potential in the non-terminating case are generated by currents close to or after the electrode location. In the terminating axon, there are no currents at points after the termination, leading to a partial suppression of the second peak and a complete suppression of the third peak.

Another generic structure found in axons is a bifurcation. To emphasize the impact of bifurcations, we simulated a single axon that bifurcates three times on each branch within a distance of 200 μm, leading to a total number of 8 collaterals leaving the bifurcation zone (Figure 1C). Note that in order to avoid confounding effects, the horizontal distances between axons in Figure 1C-E are for illustration only; all collaterals were simulated to lie on a straight line. The EFP far away from the bifurcation zone has a triphasic shape and resembles the one observed in Figure 1A, and the amplitude is proportional to the number of axon fibers. The EFP near the bifurcation zone has a biphasic shape. Although there is an initial tiny positive peak, the response is dominated by the second, negative and the third, positive peak. This waveform can again be understood by comparison to the first example (Figure 1A) containing the infinitely long axon: The tiny positive initial peak resembles the infinite case, because it is constituted by the action potential-related currents flowing within the part of the axon before the bifurcation. As the action potential passes the bifurcation zone, there are now several action potentials (one in each fiber). Because of the active nature of the action potential, the active currents are the same in each outgoing fiber as in the incoming fiber. This leads the second and third peak to be multiplied in size, yielding a quasi-biphasic response. We chose to simulate several bifurcations because this leads to a clearer effect in the EFP. In the case of a single bifurcation, this effect is also present, but the amplification of the second and third peak relative to the first peak is not as notable as in this example.

To understand how bifurcations and terminations interact when they are present in the same axon, we simulated an axon with an identical number of bifurcations as in the previous case, but then added terminations to all the fibers 200 μm after the bifurcation zone (Figure 1D). We found that this configuration leads to the same biphasic responses as observed in the cases of the isolated anatomical features. A triphasic response occurred in-between the bifurcation and termination zones. A notable point here is that the potential at the bifurcation and termination are both biphasic and on the same timescale, but opposite in polarity.

After having studied the EFPs of single axons, we next started to simulate axon bundles, because axons often run in parallel bundles in the brain. Moving towards more biologically plausible axon geometries, we considered axon bundles consisting of axons with slightly altered spatial arrangement: we randomly perturbed the precise locations of bifurcations and terminations in the axon tree (Figure 1E, left). We simulated 100 axons and stimulated each axon with an inhomogeneous Poisson spike train (Softky and Koch, 1993; Kuokkanen et al., 2010). The driving rate of the inhomogeneous Poisson process was the same for all axons and consisted of a constant background rate (100 spikes/s) and a Gaussian pulse of heightened activity (2000 spikes/s). The standard deviation of the pulse was 1 ms, resulting in an additional 3.5 spikes being fired on average over the entire duration of the pulse. The resulting extracellular population-level waveforms (Figure 1E, right) showed a polarity reversal reminiscent of Figure 1D. However, in the bifurcation zone, the summed contribution from many fibers and action potentials lead to a monophasic negative peak, and in the termination zone there was a monophasic positive peak. Interestingly, the summed potential at the center of the terminal zone largely cancelled out.

The fact that the responses in Figure 1E were mostly monophasic can be explained by the presence of a non-zero bias in the biphasic responses observed for the single spike responses in Figure 1D: close to a bifurcation, the area under the negative part of the curve slightly exceeded that of the positive part, and vice versa close to a termination. When summed up over many spikes with different timings, this difference in areas induced a positive or negative polarity of the population response in Figure 1E.

The reversal behaviour shown in Figure 1E is similar to the polarity reversal associated with a dipole observed in experimental studies (Swadlow and Gusev, 2000; Swadlow et al., 2002; Blot and Barbour, 2014). To summarize, simple one-dimensional model axon structures can produce complex and diverse spatiotemporal EFP responses, including monophasic, biphasic and triphasic waveforms, comparable to experimentally recorded responses.

### Axonal projections generate a dipole-like field potential

Dipole-like EFPs have a much larger spatial reach than quadrupolar-like EFPs, which are typically associated with axons (Nunez and Srinivasan, 2006). To further understand whether and how axons can generate a dipolar EFP, in Figure 2 we turned to three-dimensional axon morphologies, in contrast to the one-dimensional case studied in Figure 1 (for details, see Materials and Methods). We thus simulated a parallel fiber bundle of 5000 axons that at first runs at a constant number of fibers without bifurcations and then reaches a terminal zone. Within this terminal zone, the fibers first bifurcate, which increases the number of fibers. Finally, as the axons reach the end of the terminal zone, they terminate and the number of fibers decreases to zero (example axon shown in Figure 2A). To model more closely the actual axonal structures found in nature, we included a radial fanning out of the branches as well as a more diverse set of morphologies with a variable number of bifurcations and terminations (see Materials and Methods for details).

**Figure 2:**
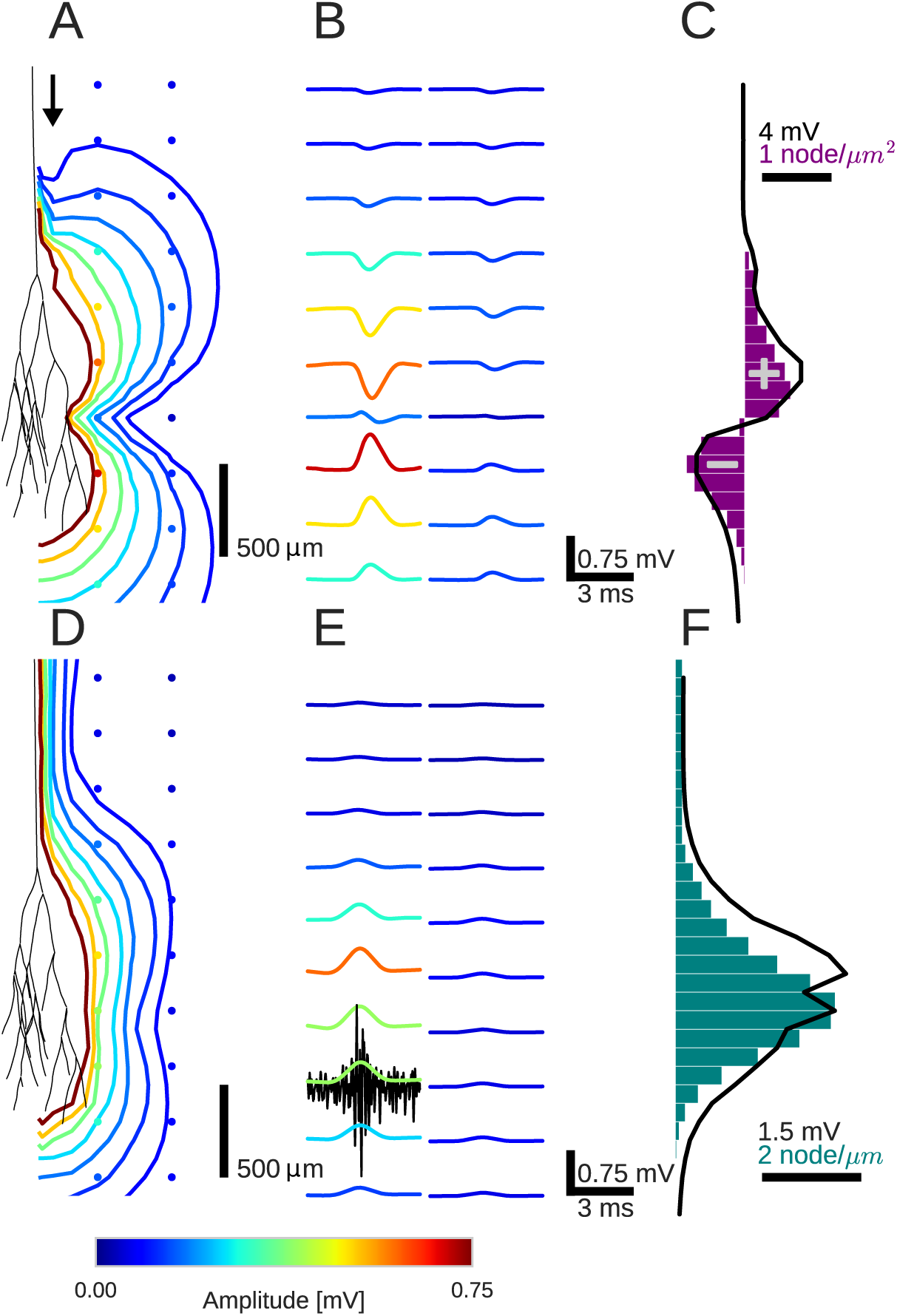
An activity pulse in an axonal projection generates a dipole-like local field potential (LFP). (**A**) Modeled example axon from the simulated bundle in black, along with iso-potential lines for the low-pass filtered (<1 kHz) EFP signature of the activity pulse. The contours (amplitudes in mV as indicated by colorbar) show the typical double-lobe of a dipole. (**B**) The LFP waveforms, recorded at the locations of the colored dots in A, show mostly unimodal peaks. The peak amplitude reverses polarity as a function of recording location in the vertical direction. The reversal occurs by inverting the amplitude with approximately unchanged shape. (**C**) Progression of the maximum LFP amplitude with depth (black line) at a distance of 100 μm from the trunk (indicated by arrow in A). The amplitude closely follows the local change in number of fibers, i.e. the difference in number between bifurcations and terminations (purple histogram). (**D**) Modeled axon from bundle as in A, and iso-potential contours for the MUA component. (**E**) Response waveforms of the MUA component. High-pass filtered (>2.5 kHz) component (the first processing stage for calculation of MUA, see Materials and Methods) in black. (**F**) Maximum amplitude of the MUA component (black line) follows the number of fibers (teal-colored histogram). Note the different units of the histograms in (C) and (F), due to the fact that (C) is the derivative in space of (F).

The spiking activity of a generic axon bundle was simulated by a background spontaneous firing rate of 100 spikes/s and a short pulse of increased activity. The responses were averaged over 10 repetitions. We chose a Gaussian pulse with an maximum instantaneous rate of 2900 spikes/s and a standard deviation of 2.8 ms. Note that this high driving rate is only the instantaneous maximum, and the actual firing rate is limited by the refractory period following a spike. These numbers are motivated by the early auditory system of barn owls (Sullivan and Konishi, 1984; Konishi et al., 1985; Köppl, 1997a), where instantaneous spike rates of 3000 spikes/s occur in response to click stimuli (Carr et al., 2016). However, our approach is not limited to the auditory system (which would also require the introduction of the synchronization of the spike times to the auditory stimulation frequency, called phase locking). Instead, this pulse of activity could relate to various kinds of evoked activity in the nervous system, such as sensory stimulation, motor activity or a spontaneous transient increase in population spiking activity.

To characterize the spatiotemporal dynamics of the evoked EFP, the time course of the potential was calculated for several locations along the axon trunk. We divided our analysis into two frequency bands by filtering the responses. The first frequency band was the local field potential (LFP) obtained by low-pass filtering with a cutoff frequency of 1 kHz (Figure 2A-C). The second frequency band was the multiunit activity (MUA) obtained by high-pass filtering with a cutoff frequency of 2.5 kHz (Figure 2D-F). To make the MUA easier to interpret in terms of overall activity reflected, it was half-wave rectified and low-pass filtered (<500 Hz, see Materials and Methods). The two frequency bands showed a qualitatively different spatiotemporal response in the vicinity of the projection zone, as we will show in the following.

We first studied the effect of the Gaussian rate pulse on the LFP (Figure 2B). The filtering used to obtain the LFP removed most of the identifiable components of individual spikes, while a population-level LFP signal remained. The distribution of the maximum amplitudes of these responses is shown by the colored contour lines in Figure 2A and the colored voltage traces in Figure 2B. Surrounding the terminal zone of the axon bundle in Figure 2A, LFP amplitudes showed a double-lobed shape typical of a dipole.

In Figure 2B, the LFP responses mostly showed monophasic deflections elicited by the population firing rate pulse, in a manner similar to that observed in Figure 1E. Such deflections were visible at all recording locations. In the radial direction away from the axon tree, i.e. in the horizontal direction in the figure, the LFP amplitude decays. In the axial direction along the axon tree, i.e. in vertical direction in the figure, the voltage deflection reverses polarity in the middle of the terminal zone of the bundle (Figure 2B). The polarity reversal occurs by a decrease of the amplitude to zero and a subsequent reappearance with reversed polarity (as opposed to a polarity reversal through a gradual shift in phase). This behaviour is also typical for a dipolar field potential.

The point of the polarity reversal coincides with the middle of the terminal zone. Interestingly, this means that the absolute value of the response amplitude reaches a local minimum just when the number of axonal fibers reaches a maximum. To better understand how the anatomical features of the axon bundle and the LFP response amplitude are related, we compared its signed maximum value (meaning the ***signed*** value corresponding to the maximum *magnitude* of the amplitude of the LFP, black line in Figure 2C) with the difference between the number of branchings and terminations per 200 μm bin (purple histogram in Figure 2C): Along the nerve trunk the number of fibers is constant. As the axon bundle reaches its terminal zone, the number of bifurcations increases (purple bars point to the right in Figure 2C). The increase of bifurcations is followed by an increase in terminations. In the middle of the terminal zone, the number of bifurcations and terminations are equal. At the same depth, the amplitude of the LFP component crosses zero. At the end of the terminal zone, the terminations outweigh the bifurcations (purple bars point leftwards in Figure 2C). As the axon bundle ends, there are no longer any bifurcations or terminations, and the number of fibers decays toward zero. Overall, the signed maximum amplitude LFP (black trace) follows the distribution of branchings and terminations (purple histogram). This progression of amplitudes in the low-frequency components seen in Figure 2C is also visible in Figure 2B, most clearly in the first column. The polarity reversal in the center of Figure 2B corresponds to the crossing of zero amplitude in Figure 2C.

To understand how the EFP contributions are related to individual spikes, we next turned our attention to the high-frequency MUA response. The MUA is thought to reflect local spiking activity (Stark and Abeles, 2007). In Figure 2D, the iso-amplitude lines of the MUA appeared like an ellipsoid centered at the terminal zone (Figure 2D); they did not show the double-lobe shape observed for the LFP in Figure 2A.

The shape of the MUA response was weakly dependent on the recording location. The main change across locations was in the scaling of the amplitude (Figure 2E). The amplitude decays with radial distance from the trunk. In the axial direction, the amplitude reaches its maximum in the middle of the fiber bundle. This dependence of the MUA amplitude on the axial location is further examined in Figure 2F. The amplitude of the MUA component (black trace) changes in accordance with the local number of fibers (teal-colored histogram). The local number of fibers and the MUA amplitude are both constant along the nerve trunk. Both measures then increase in amplitude as the number of fibers is increased by bifurcations. As the fibers terminate and the number of fibers decreases, so does the amplitude of the MUA.

To conclude, we have shown a qualitatively different behaviour in the low- and high-frequency components of the EFP, i.e. for the LFP and the MUA. The particular branching and terminating structure of the axon bundle may thus give rise to a dipolar LFP.

### Effects of bifurcations and terminations on distance scaling of EFPs

To further demonstrate that bifurcations and terminations of axons give rise to a dipolar field, we investigated the effect of an axon terminal structure on the spatial reach of the EFP (Figure 3). Motivated by the fundamentally different spatial distributions of the low-frequency LFP and the high-frequency MUA in Figure 2, we again differentiated between these frequency bands and simulated an axon bundle containing a terminal zone with bifurcations and terminations. Moreover, as a control, we also simulated an axon bundle without bifurcations in which a fixed number of fibers simply terminates.

**Figure 3:**
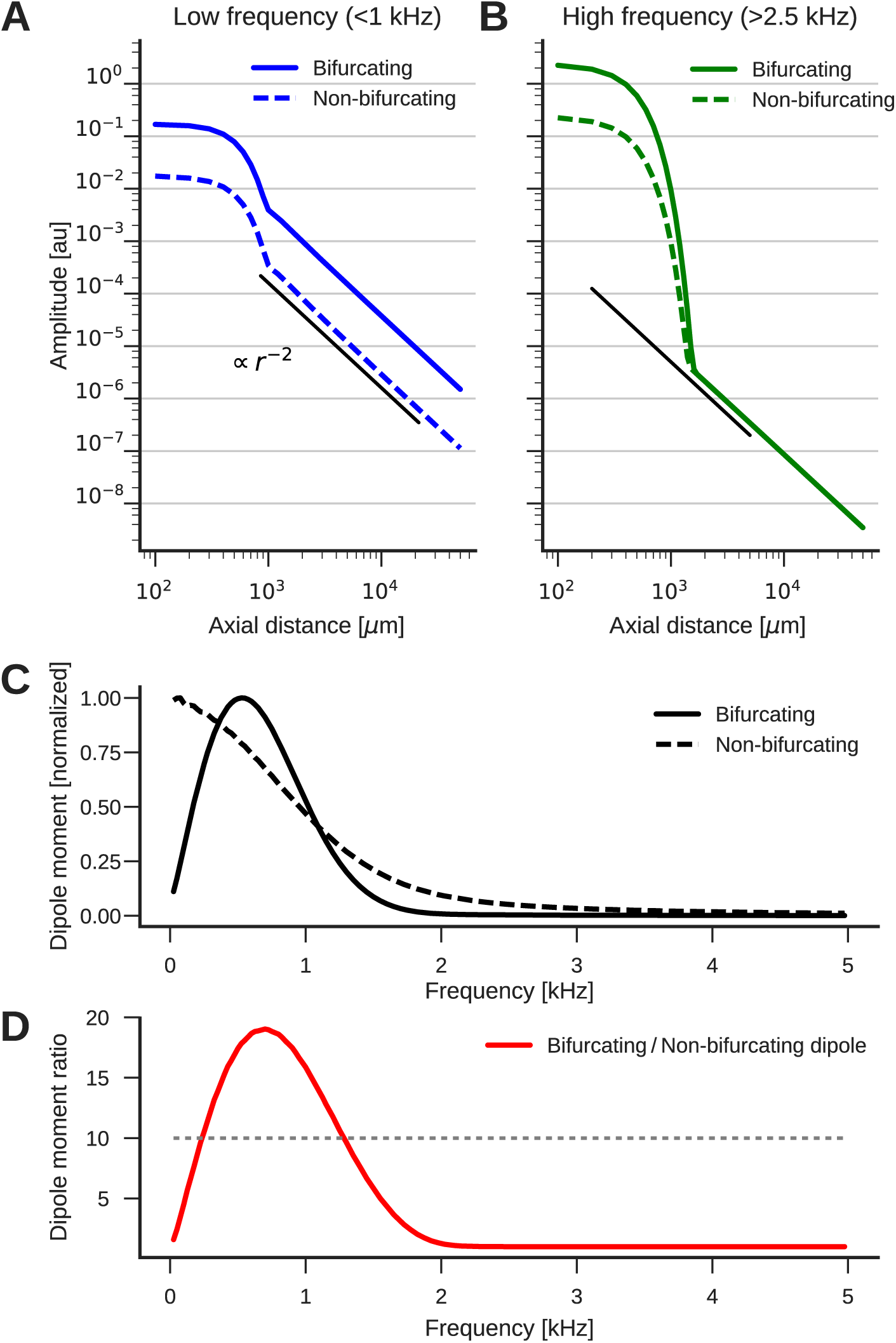
The low-frequency (<1 kHz) component of the axon bundle EFP exceeds the high-frequency (>2.5 kHz) component in reach. The scaling of the low-frequency (**A**, blue lines) and high-frequency (**B**, green lines) components for the bifurcating (solid lines) and non-bifurcating (dashed lines) axons on a double-logarithmic scale. All distances are calculated from the center of the terminal zone. For comparison, scalings that follow *r*^−2^ are shown (black lines). (**C**) Normalized dipole moments of the bifurcating and non-bifurcating bundles as a function of frequency. (**D**) Ratio of the dipole moments between bifurcating and non-bifurcating cases (red line), compared to the maximum ratio 10 of the number of fibers (dotted line), to indicate supralinear (>10) and sublinear (<10) contributions.

In order to separate the effects of any radial fanning out of the axon bundle from the effects of bifurcations and terminations, and to afford better analytic tractability, we transitioned back to a simpler one-dimensional model of the axon bundle (see Materials and Methods). This model omitted the radial fanning out of the bundle in the terminal zone, as in Figure 1. Furthermore, we discarded the detailed conductance-based simulation of the membrane potential, and instead assumed a fixed membrane potential waveform traveling along the axon trunk with a constant propagation velocity. Using linear cable theory, it was then possible to calculate the membrane currents necessary for the determination of the EFP. The analytic nature of the simplified model also allowed us to consider a continuous number of fibers instead of simulating discrete bifurcations and terminations. All following simulations are based on this simplified model.

We simulated two axon bundle morphologies. The control case was a non-bifurcating bundle, which had a constant number of 50 fibers up to the termination point, and then tapered out with a Gaussian profile that was centered at the termination point with a height of 50 fibers and width of 300 μm. The second case was that of an axon bundle with a projection zone containing bifurcations. Here we added to the distribution of the number of axons used for the non-bifurcating control case a further Gaussian distribution to account for the projection zone. This additional Gaussian was also centered at the termination point, but had a width of 500 μm and an amplitude of 450 fibers. Unlike the tapering-out in the control condition, this component was added for both before and after the termination point. It resulted in a maximal fiber number of 500 at the termination point, which is a factor 10 larger than in the control case. Both distributions constructed in this way were smooth, and they had smooth first derivatives in space. We considered a conduction velocity of 1 m/s in this example, though results are qualitatively the same for other values. In order to understand the frequency-specific effects of the projection zone, we calculated the responses to membrane potential components with temporal frequencies between 100 Hz and 5 kHz, with the same amplitude for each frequency component. Due to the linear nature of our model, the frequency responses obtained in this manner are applicable to the Fourier components of any membrane potential time-course. We then calculated the average amplitude of the resulting EFP response by taking the standard deviation.

The dipole-like component observed in Figure 2 for the low-frequency component had its dipole axis aligned with the axon trunk. We therefore considered the distance *r* beyond the termination point in the direction extending the axon trunk, which we call the axial direction. Because we suspected a dipolar response, we expected the amplitude of the field potential to decay as *r*^−2^. To test the scaling behaviour of this component, we first plotted the average amplitude of the low-frequency responses (*f* <1 kHz) in axial direction in Figure 3A. The plot is on a double logarithmic scale, meaning that the slope of the curve corresponds to the scaling exponent, and the vertical offset corresponds to the amplitude of the size of the dipole moment, which is a measure for the strength of the dipolar EFP. We observed the expected *r*^−2^ scaling for distances larger than the extent of the bifurcation zone (≳ 1 mm).

Comparing the responses of the bifurcating axon bundle and the non-bifurcating control condition (full and dashed lines in Figure 3A), we saw that for short distances (< 1 mm) the response of the bifurcating case was a factor 10 larger than the control. At these distances the response was due to the local fibers, of which there are 10 times more in the bifurcating case. At distances larger than 1 mm, we observed that the distance scaling was proportional to *r*^−2^, meaning that there was a dipole moment in both conditions (a vanishing dipole moment would have implied a slope steeper than *r*^−2^). Interestingly, for distances larger than 1 mm the response in the bifurcating case exceeded the control by a factor 20. We thus concluded that at low frequencies, the bifurcation zone contributes supralinearly to the dipole moment.

Next, we examined the high-frequency (>2.5 kHz) component (Figure 3B). As in the low-frequency case, the response at distances < 1 mm was greater in the bifurcating case by a factor of 10. The asymptotic scaling was also *r*^−2^ for axial distances > 1 mm in both cases. However, unlike in the low-frequency case, the amplitudes were similar between bifurcating and non-bifurcating cases. This meant that the presence of a bifurcation zone did not contribute to the high-frequency dipole moments in the EFP.

### Frequency-dependence of dipolar axonal EFPs

We showed that the dipole moment depends on both the anatomy, i.e. the presence of a projection zone, and the temporal frequency range of the underlying activity. This relationship can be qualitatively understood by considering that in an axon bundle a voltage waveform propagates at some conduction velocity.

The temporal frequency of this signal thus corresponds to a spatial frequency. If the spatial frequency of the membrane potential matches the width of the projection zone, the dipole moment can reach its maximum. In this case, at some point in time, positive membrane currents flow in one half of the projection zone and negative membrane currents flow in the other half. For example, if the voltage waveform has a temporal frequency of 1 kHz and propagates at 1 m/s, the spatial wavelength is 1 mm. If the spatial width of the termination zone is about 1 mm, the dipole moment is maximal (for a detailed derivation see Materials and Methods).

To quantitatively understand the frequency-specific contributions to the dipole moments, we examined the scaling behaviour of the EFP as a function of frequency. The amplitude of the dipole moment was determined by fitting a straight line with slope –2 to the double logarithmic scaling of the standard deviation of the response at a given frequency. The fit was performed for distances > 1 mm. The extrapolation of this straight line to the axial distance 1 μm was then proportional to the dipole moments.

The normalized frequency-specific dipole moments are shown in Figure 3C. The dipole moment of the bifurcating case (solid line) has a maximum at around 750 Hz, as expected due to the agreement of the spatial wavelength (1 m/s / 750 Hz = 1.33 mm) and axial width (about 1 mm) of the projection zone. The match is not exact because the shapes of sine wave and Gaussian are different. For lower and higher frequencies, there is a mismatch in spatial wavelength and the width of the projection zone, meaning that the projection zone contributes only little to the dipole moment, as observed in Figure 3B for higher frequencies. In the non-bifurcating control case (dashed line) there is no projection zone, and the dipole moment decays monotonically because lower frequencies correspond to a larger spatial separation of positive and negative currents, and thus to a higher dipole moment.

In the bifurcating case, the maximum number of fibers was increased by a factor of 10. Accordingly, an increase in the dipole moment by a factor of 10 from the non-bifurcating to the bifurcating case could be explained by just linearly summing the dipole moments of individual fibers. An increase in the dipole moment by a factor greater than 10 would be supralinear. In Figure 3D we compared this relative impact of the terminal zone on dipole moments (red line) by plotting the dipole moment ratios across frequencies. The contribution of the terminal zone is greater than 10 (dotted line) for intermediate frequencies between about 200 and 1300 Hz, and smaller than a factor 10 outside this frequency range.

Together, these observations show us that the terminal zone makes a frequency specific contribution to the far-reaching dipole field potential of the axon bundle. This provides a deeper understanding of the findings of Figure 2: At low frequencies (< 1 kHz), we observed a supralinear dipolar behaviour due to the strong contribution of the bundle at these frequencies, with the projection zone forming the dipole axis. At higher frequencies, the bifurcation zone strongly reduces the dipole moment, meaning that we could observe responses mainly locally.

### The barn owl neurophonic potential in nucleus laminaris as an example for a dipolar field in an axonal terminal zone

To test our models prediction of dipolar extracellular field potential responses due to axon bundles, we recorded EFP responses from the barn owl auditory brain stem. The barn owl has a highly developed auditory system with a strong frequency-following response in the EFP (up to 9 kHz, Köppl (1997b)), called the neurophonic, which can be recorded in the nucleus laminaris (NL). In NL, the input from the two ears is first integrated to calculate the azimuthal location of a sound source, and this information is encoded in the EFP (Carr and Konishi, 1990). The EFP in this region is mainly due to the afferent activity, and the contribution of postsynaptic NL spikes is small (Kuokkanen et al., 2010, 2013). Furthermore, the anatomy of the afferent axons is well known and follows a stereotypical pattern (Carr and Konishi, 1988, 1990): Two fiber bundles enter the nucleus, with fibers from the contralateral ear entering ventrally, and from the ipsilateral ear entering dorsally. The axon bundles reach the NL from their origin without bifurcating, then bifurcate multiple times at the border of the NL, and then terminate within NL. Axon bundles have a strong directional preference and run roughly in parallel. Most of the volume within NL consists of incoming axons. This well studied physiology and anatomy makes the system an ideal candidate to investigate the EFPs of axon bundles; see the Discussion for arguments why synaptic contributions to the EFP could also be neglected here.

To explore the spatiotemporal structure of the EFP in NL, we performed simultaneous multielectrode recordings of the response in NL (Figure 4A) to contralateral monaural click stimuli. The click responses showed distinct low-frequency (Figure 4B) and high-frequency (Figure 4C) components, as previously reported (Wagner et al., 2009). The frequency of the high-frequency ringing corresponds to the recording location on the frequency map within NL, and the ringing reflects the frequency tuning and phase locking of the incoming axons. In addition, there is a low-frequency component in the response (Figure 4B). We filtered the data to roughly separate these components. The cutoff to split the components was set to 2 kHz because this frequency was always well below the high-frequency ringing component.

**Figure 4:**
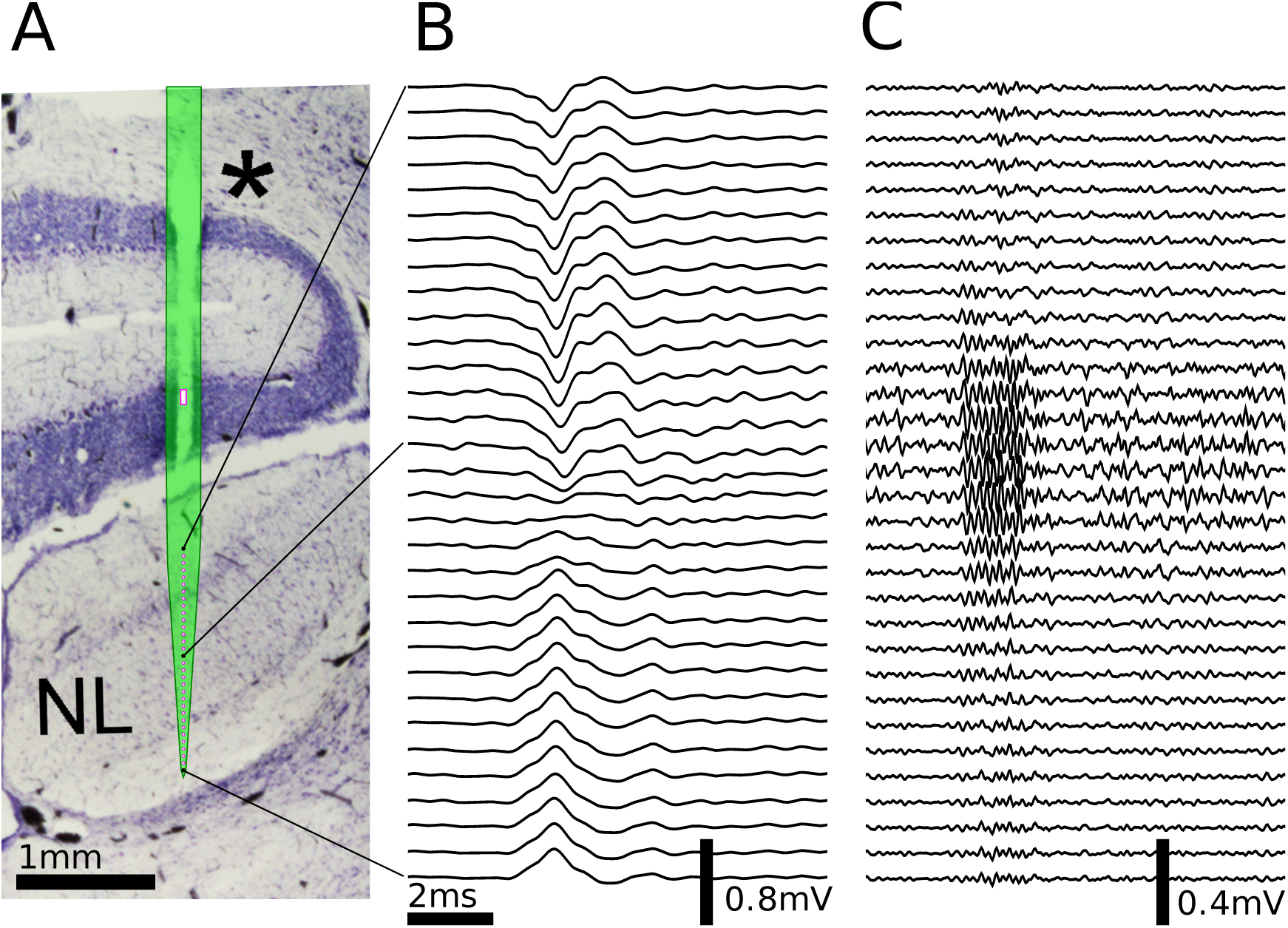
Multielectrode recordings in the barn owl show dipolar axonal EFPs. (**A**) Photomicrograph of a 40 μm thick transverse Nissl stained section through the dorsal brain stem, containing a superimposed, to scale, diagram of the multielectrode probe. The probe produced a small slit in a cerebellar folium overlying the IVth ventricle (*), and penetrated into the nucleus laminaris (NL). The recordings were made in NL, and electrodes extended to both sides of the nucleus. The outline of the probe is shown in light green, with the recording electrodes indicated by magenta dots, and the reference electrode as a magenta rectangle. The low-frequency (<2 kHz) component (**B**) and the high-frequency (>2 kHz) component (**C**) are ordered in the same way as the electrodes, with three examples connected to their recording sites by black lines. The time scales in B and C are identical (indicated by scale bar). Traces were averaged over 10 repetitions. Voltage scales are indicated by individual scalebars.

The same simplified model used in Figure 3 was fit to the data (example in Figure 4) by performing a nonlinear least squares optimization. Free parameters were (1) the number of fibers at the depth of each recording location, (2) the average spatial derivative of the membrane potential over time in the fibers at the location next to the most dorsal electrode, (3) the axonal conduction velocity, and (4) the distance between the axon bundle and electrode array.

The resulting EFP responses and the model fit are depicted in Figure 5. Figure 5A shows the inferred average over trials of the deviation of the membrane potential from the resting potential in response to the stimulus, at a location in the axon next to the first electrode (penetration depth 1550 μm), obtained from the fit. The inferred voltage is composed of high- and low-frequency components similar to those observed in the EFP. The inferred number of fibers as a function of dorsoventral depth is shown in Figure 5B. The number (scaled by an arbitrary factor) has its maximum at the center of the electrode array, and decays steadily to both sides. This profile of the number of fibers is consistent with the known anatomy of axons in NL (Carr and Konishi, 1990; Kuokkanen et al., 2010).

**Figure 5:**
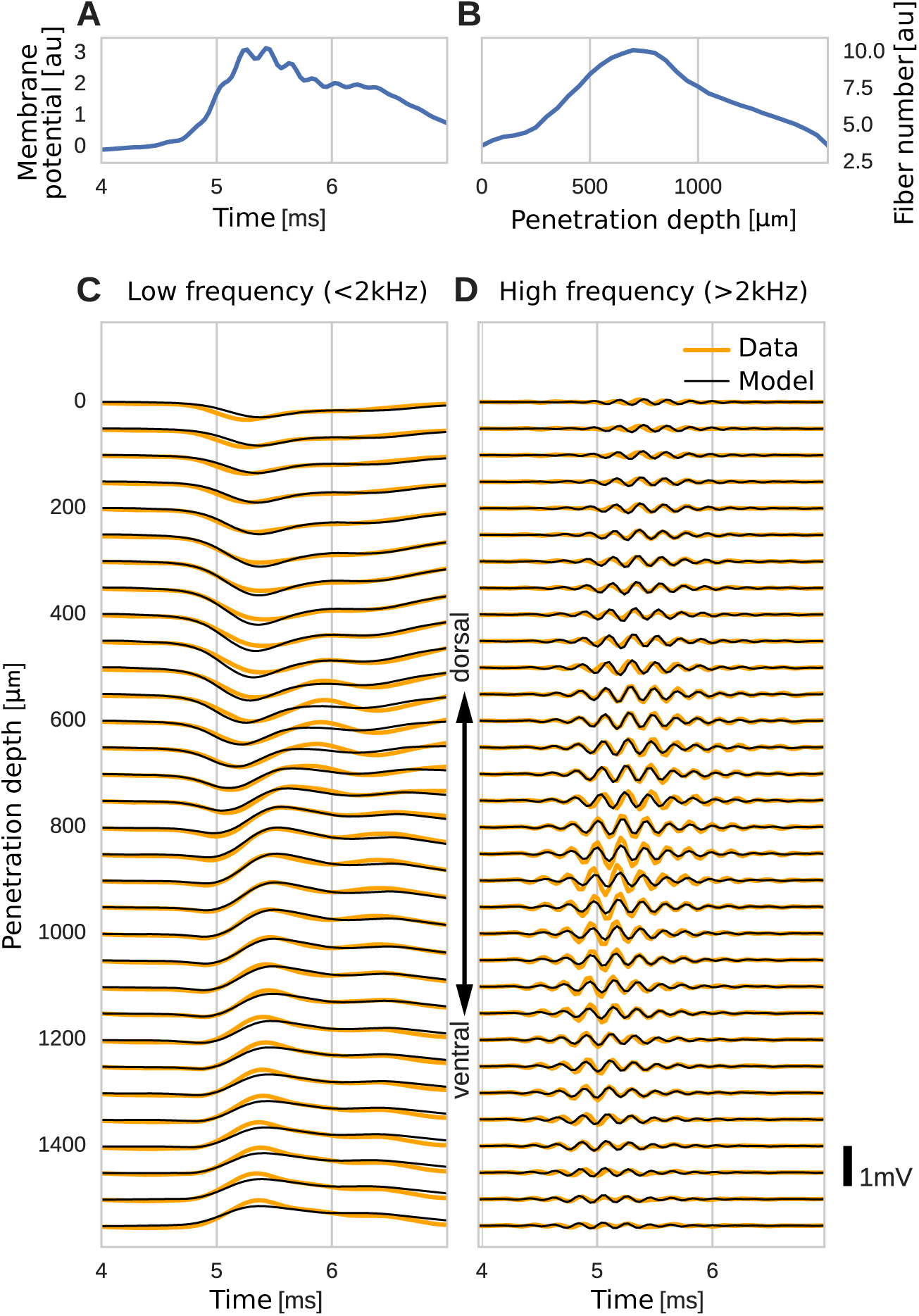
The spatial structure of EFPs recorded from the nucleus laminaris of the barn owl can be explained by a model of axonal field potentials (for details, see Materials and Methods). (**A**) Membrane voltage averaged across fibers in the model when fit to the data. (**B**) Fitted number of fibers in the model as a function of penetration depth. (C) Low-frequency (< 2 kHz) components of the EFP in response to a click stimulus at time 0 ms, at different recording depths. The depth is measured in the direction from dorsal to ventral. Recorded responses (orange) are shown along with model fits (black). (**D**) High-frequency (> 2 kHz) responses in recordings (orange) and model (black). Recorded traces were averaged over 10 repetitions.

The low-frequency (30 Hz - 2 kHz, Figure 5C) responses reveal the typical polarity reversal that we predicted for an axonal terminal zone (Figures 1, 2). The dorsoventral depth is now on the vertical axis, meaning that the vertical axis of Figure 5C and D corresponds to the horizontal axis in Figure 5B. The orange lines indicate the actual responses in the data.

The responses at the dorsal and ventral edges show the same shape, but with opposite polarity, as expected for a dipolar field. Note that for a pure dipole field, the amplitude of the central responses have zero amplitude. In the data shown here, central responses show a diminished maximum amplitude, which we interpret as the contribution of higher-order (mostly quadrupole) components. The model is able to capture the behaviour of this quadrupolar component as well, with a slight underestimation of the amplitude of the peak at ventral locations. The model even captures a small oscillation in the data with period of ≈ 1 ms in the center of the recording. Here, too, the small deviations are likely due to slightly inhomogeneous conduction velocities or non-axonal sources.

In addition to the dipolar behavior of the low-frequency response, we also examined the high-frequency (≥ 2 kHz) response, shown in Figure 5C. The responses have a Gabor-like shape, as expected (Wagner et al., 2009), with maximum amplitude in the center of the recording array, at around 850 μm penetration depth. The axonal conduction velocity was calculated to be 4.0 m/s, and the distance from the bundle was 162 μm. A previously published estimate of the axonal conduction velocity in this nucleus (McColgan et al., 2014) gave a confidence bound of 0.4-6 m/s. Toward the edges (< 100 μm and > 1400 μm), the amplitude decays. In the central region (400-1200 μm recording depth), a systematic shift in delay can be observed, while the response appears stationary in the more dorsal and ventral electrodes. The delay increases from ventral to dorsal, which is consistent with the anatomy for contralateral stimulation.

All these aspects of the data are qualitatively reproduced by the model (Figure 5D, black traces). The main deviation between model and data lies in a diminished amplitude of the oscillation modelled at the most central electrode sites. Because the phase shift in the central region is mainly determined by the conduction velocity, this mismatch might be due to a variable conduction velocity in the nucleus, and the constant velocity in the model. McColgan et al. (2014) showed that different conduction velocities exist in the core and periphery of the nucleus, as predicted from variable internode distances by Carr and Konishi (1990). A diminished amplitude in the fit could reflect an inability of the model to exactly match the phase progression. Another possible explanation is that the additional amplitude could be due to non-axonal sources such as synaptic currents or postsynaptic spikes, which do not follow the assumptions underlying our model; see the Discussion for arguments why we expect such contributions to the EFP to be small.

When comparing the inferred membrane potential response (Figure 5A) to the measured EFP response (Figure 5C and D), the most salient difference is the dissimilar sizes of the frequency components. In the EFP, the low-frequency component has a comparable amplitude to the high-frequency component, but in the membrane potential the low-frequency component is much larger because the EFP is related to membrane currents, which are proportional to the first and second derivatives of the membrane potential, and taking the derivative is equivalent to applying a high-pass filter.

We performed the fitting procedure (example in Figure 5) for 26 recordings from 3 different owls, with monaural stimulation from both ears (implying the activation of distinct axonal populations). The average correlation coefficient for all recordings was *R*^2^ ***=*** 0.56 ± 0.15. The correlation coefficient for the example shown in Figure 5 was 0.62.

### Dipole moments of idealized axon bundles

We have shown theoretically and experimentally for specific examples of axonal projection zones and inputs how dipolar EFPs emerge. We now generalize this approach and predict the resulting dipolar EFP for arbitrary axon and stimulus configurations. Based on our cable-theory model, we analytically derived the maximal dipole moment *p*_max_ for a large range of scenarios. From a given dipole moment the maximum far field potential at distance *r* can be calculated as 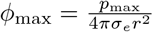 where *σ_e_* is the extracellular conductivity.

To simplify the analytical derivation as much as possible, we assumed a Gaussian waveform for the membrane potential of a single spike, with an amplitude *V̅*_spike_ and a width σ_spike_. The resting membrane potential was irrelevant because only the first and second derivatives of the membrane potential contribute. The axon bundle population consisted of fibers with diameter *a*, axial resistance *r_L_,* and conduction velocity υ. The population was assumed to be driven with a Gaussian firing-rate pulse with maximum firing rate λ̄_pulse_ and width σ_pulse_. The distribution of the number of fibers at a given depth location was also described with a Gaussian, with width *σ*_***n***_ and maximum number *n̄*. This is an adequate approximation if the spikes in the incoming fibers contribute little to the dipole moment before reaching the projection zone. In this scenario, we calculated the maximum dipole moment of the bundle (see Materials and Methods for details) to be

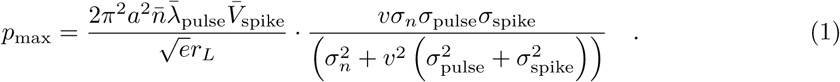

Equation 1 tells us that the dipole moment is proportional to *a*^2^, *n̄*, λ̄_pulse_, *V̅*_spike_, and 1/*r*_L_. The dependence on *v* and the widths is more complicated; the response is maximal with respect to the three (spatial) widths σ_*n*_, *vσ*_pulse_ and *vσ*_spike_ when they are of the form *w*_1_^2^ = *w*_2_^2^ + *w*_3_^2^, where *w*_1_ is the largest of the three terms, while *w*_2_ and *w*_3_ are the other two terms, regardless of order. The dipole moment is thus maximal when the widths of the spike, the pulse, and the terminal zone agree. In particular, if *σ_n_* (the width of the terminal zone) is the widest, then the dipole moment is maximal if *σ_n_* is equal to the spatial width of the overall activity in the axons, which is 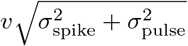. The widths add in this way because the overall activity is the convolution of two Gaussians.

Using this formula, it is then possible to calculate the expected contributions to the EFP for different scenarios. To test the approximation in the case of the barn owl, we chose the following values: axon diameter *a* = 2 μm, conduction velocity 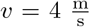 as inferred in the previous section, axial resistivity *r_L_* = 1 Ωm, and extracellular conductivity 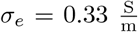 as used in studies of the cortex (Holt and Koch, 1999; Gold et al., 2006), anatomical and physiological parameters *σ_n_* = 500 μm, *n̄* = 4000, *V̅* S_pike_ = 70 mV from (Carr and Konishi, 1990), and activation patterns for click stimulation from (Köppl, 1997a; Carr et al., 2016): λ̄_pulse_ = 3000 spikes/s, σ_spike_ = 250 μs, σ_pulse_ = 0.5 ms. This leads to a value for the dipole moment of *p*_max_ ≈ 1.9 μA · mm. At a distance of 750 μm, roughly the furthest distance recorded with the multielectrode array in Figure 4 and Figure 5, this dipole moment corresponded to a field potential of 0.82 mV, consistent with our experimental findings.

Dipole sources are also to be expected to make up the majority of the electrical signals recorded at the scalp (Nunez and Srinivasan, 2006). One such signal is the auditory brainstem response (ABR), which is recorded at the scalp in response to auditory stimulation with clicks or chirps (Riedel and Kollmeier, 2002). The amplitude of about 10 μV of the ABR in the barn owl has recently been reported by Palanca-Castan et al. (2016). We calculated the contribution expected from an axon bundle with the same characteristics as described before at 2 cm from NL, aiming to estimate the contribution to the ABR. Multiplying by a factor of 2 to account for the fact that there is an NL in each hemisphere, the predicted contribution was 2.3 μV, which is of the same order of magnitude as the value reported in the experiments.

As a second example, we considered thalamocortical projections, for which Swadlow and Gusev (2000) reported amplitudes of extracellular spike-related potentials, called axon terminal potentials (AxTP), at various locations; for example, at 400 μm from the center of the dipole, they reported an amplitude of the response of ≈ 1 μV. Individual thalamocortical axons are thin and have large and highly branched projection zones (Feldmeyer, 2012), so we estimated *σ_n_* = 250 μm, *n̄* = 30, and *a* =1 μm. We assumed a jitter *σ*_pulse_ = 125 μs in the arrival time instead of a true activity pulse, and we normalized the pulse to have area 1 because we were considering a spike triggered average. The conduction velocity has been reported as *v* = 8.5 m/s (Simons et al., 2007). Leaving all other values as in the previous approximation, we arrived at a dipole moment of *p*_max_ ≈ 1.5μA · μm, yielding an extracellular spike amplitude of ≈ 2.3 μV at the distance of 400 μm, which is of the same order of magnitude as the value (≈ 1 μV) reported by Swadlow and Gusev (2000).

To summarize, Equation 1 quantitatively predicts the contribution of axonal projection zones to the far field EFP, and this prediction matched experimental values in several cases.

## Discussion

Numerical simulations, analytical calculations, and experimental data allow us to show how axonal fiber bundles may contribute to the EFP, and explain how the contributions are shaped by axonal morphology. There are three principal effects of axon bundle structure on the EFP. First, the low-frequency components of the EFP are governed by the densities of bifurcations and terminations and can have a dipolar structure (Figure 1 and Figure 2A-C). Second, the high-frequency components are governed by the local number of fibers (Figure 2D-F). Third, the low-frequency components exceed the high-frequency components in spatial reach. In particular, the dipolar low-frequency components are not negligible and exceed the reach of the presumed quadrupolar nature of axonal EFPs (Figure 3).

### Relevance to the interpretation of electrophysiological recordings

Our findings relate to the interpretation of a wide range of electrophysiological data in general, and to the estimation of current sources in particular. When performing a typical current source density (CSD) analysis, the local number of fibers cannot be disentangled from membrane current density (Nicholson, 1973; Potworowski et al., 2011). In CSD analysis, the membrane current densities can independently vary with time and location. In the case of an axonal fiber bundle as discussed here, the situation is different: the number of fibers is variable in space, in particular in the terminal zone, but the current sources at different locations are highly determined, because they are caused by propagating action potentials. In the case presented here (Figure 5), where axonal action potentials dominate the EFP, it was possible to recover from the recording the (normalized) fiber densities and average membrane potentials.

Beyond recovering actual fiber densities and membrane potentials, our approach enables the interpretation of CSD results in the presence of axon fiber bundles. For example, the sink and source distribution found in classical CSD analysis of axon bundles (Mitzdorf and Singer, 1977, 1978; Mitzdorf, 1985) shows a dipolar structure in terminal zones, but a conclusive explanation of their origin was not given. Our modeling approach provides a novel way of interpreting these findings in terms of actively propagated action potentials in a fiber bundle.

As an example case for a fiber bundle, we recorded from the barn owl nucleus laminaris. Figure 4 and Figure 5 showed that the low- and high-frequency components show qualitatively different behaviours as a function of recording location relative to the terminal zone. The low-frequency component is a largely stationary phenomenon, while the fine structure of the high-frequency component shifts gradually in space as a function of the axonal conduction velocity (Figure 5). Low-frequency components have a strong dipole moment, meaning that it contributes to the far-field EFP. Due to the difference in reach, the high-frequency component is most suitable for the study of local phenomena while the low-frequency component bears information about locations more distant from the recording site (Figure 3), consistent with findings on non-axonal EFPs (Pettersen and Einevoll, 2008; Łęski et al., 2013).

Dipolar fields are essential for the generation of electrical field potentials at greater distances from the brain. The most prominent of these is the EEG, which is commonly attributed to the dipolar contributions of pyramidal cells (Nunez and Srinivasan, 2006). We propose that axonal contributions might also be relevant in the analysis of these fields. This is particularly true for the auditory brainstem response (ABR), which is closely related to the EEG and involves brain structures that display high degrees of synchrony as well as axonal organization, and are thus ideal candidates for the generation of axonal field potentials visible at long ranges. This would in turn have implications for the interpretation of the ABR in clinical contexts.

The ABR of the barn owl has been reported to be on the order of 10 μV (Palanca-Castan et al., 2016) while we estimated a contribution of about 2 μV amplitude from the incoming axons in NL alone. This estimate of 20% axonal contribution to the ABR suggests that there may indeed be measurable components due to axons in the ABR, in particular, and the EEG, in general. However, this estimate is crude because it did not take into account the anatomy of the skull except for its size. Future studies based on a more detailed skull model and paired recordings of ABR and EFP should improve our understanding of axonal contributions to the ABR.

We have shown that the EFP in the barn owl NL is consistent with a model of axonal sources. We believe synaptic contributions to be small in this case for the following reasons: Deflection of the somatic membrane potential due to synaptic currents is much smaller than the impact of postsynaptic spikes (Ashida et al., 2007; Funabiki et al., 2011). Since postsynaptic spikes contribute little to the EFP (Kuokkanen et al., 2010), we suspect that the synaptic contributions to the EFP are also small. Furthermore, synaptic EFP contributions would require a spatial separation of currents, which is not possible to achieve in NL because of the symmetrical arrangement of synapses on the spherical NL cell bodies (Carr and Konishi, 1990), meaning that synaptic sources can also not explain a dipolar EFP, and are thus expected to contribute little to the EFP.

### Dipolar EFPs in other animals and brain regions

It is interesting to note that a similar dipole-like reversal of polarity as shown here for the barn owl NL has been reported in the chicken NL (Schwarz, 1992), as well as in the mammalian analog to NL, the medial superior olive (MSO) (Mc Laughlin et al., 2010). The phase reversal in this case was modeled based on the assumption that the postsynaptic NL and MSO dendrites with their bipolar morphology generate the dipolar EFP response (Mc Laughlin et al., 2010; Goldwyn et al., 2014). However, in the owl this dipolar morphology of neurons is largely absent (Carr and Konishi, 1990), making dendritic sources unlikely. This suggests that similar dipolar field potentials in owls and mammals emerge from different physiological substrates. Such a convergence might point towards an evolutionary pressure favouring such an EFP structure in coincidence detection systems, and indeed, Goldwyn and Rinzel (2016) have proposed a model in which this extracellular potential enhances the function of coincidence detectors through a form of ephaptic interaction. Their approach centers on dendrites and is not directly transferable, but it seems possible that a similar mechanism might arise in the barn owl NL based on axonal EFPs.

The key assumption underlying our modeling of axonal geometries is that there exists a preferential direction of the axon arbor. In many structures this is the case, for example in the parts of the auditory brainstem we studied here. In other brain regions, this tendency is not as prominent, with a spectrum existing between completely directed and undirected growth. More undirected growth would lead to a more diffuse response in the EFP, and eventually to a cancellation of the dipolar field potentials. Cuntz et al. (2010) and Budd et al. (2010) studied the principles underlying the growth patterns of axons and found that the degree of direction in the growth of an axon depends on the balance struck between conduction delay and wiring cost. Optimizing for minimal conduction time leads to highly directed structures, while optimizing wiring cost leads to more tortuous, undirected growth. This suggests that directed structures - and thus also strong, dipolar EFPs due to axons - may be more prevalent in systems which require high temporal precision in the information processing.

### Conclusion

Axonal projections can contribute substantially to EFPs. Our results quantitatively show how the anatomy of axon terminal zones and the activity in axons determine its frequency-specific far-field contribution to the EFP.

## Materials and Methods

### Experimental recordings

The experiments were conducted at the Department of Biology of the University Maryland. Data was collected from three barn owls *(Tyto furcata pratincola).* Procedures conformed to NIH Guidelines for Animal Research and were approved by the Animal Care and Use Committee of the University of Maryland. Anaesthesia was induced prior to each experiment by intramuscular injection of a total of 8 – 10 ml/kg of 20% urethane divided into three to four injections over the course of 3 hours. Body temperature was maintained at 39°C by a feedback-controlled heating blanket.

Recordings were made in a sound-attenuating chamber (Acoustic Systems Inc., Austin, TX, USA). Tungsten electrodes with impedances between 2 and 20 MΩ were used to find suitable recording locations in NL. Once NL had been located, the tungsten electrode was retracted and replaced with a 32 channel multi-electrode array (A1×32-15mm-50-413-A32, Neuronexus, Ann Arbor, MI, USA). The multi-electrode array was lowered using a microdrive (MP225, Sutter Instruments Co., Novato, CA, USA) during continuous presentation of a white-noise burst stimulus until visual inspection of the waveform showed that NL was at the center of the array. A grounded silver chloride pellet, placed under the animal’s skin around the incision, served as the animal ground electrode. Electrode signals were amplified by a headstage (HS36, Neuralynx Inc., Tucson, AZ, USA). An adapter (ADPT-HS36-N2T-32A, Neuralynx Inc.) was used between the electrode and the headstage. A further adapter (ADPT-HS-36-ERP-27, Neuralynx Inc.) was used between the headstage and the control panel in order to map all 32 channels to the amplifiers. Pre-amplified electrode signals were passed to the control panel (ERP27, Neuralynx Inc.), then to four 8-channel amplifiers (Lynx-8, Neuralynx Inc.), and then to an analogue-to-digital converter (Cheetah Digital Interface, Neuralynx Inc.) connected to a personal computer running Cheetah5 software (Neuralynx Inc.) where the responses were stored for off-line analysis.

Acoustic stimuli were digitally generated by a custom-made MATLAB (MathWorks, Natick, MA, USA) script driving a signal-processing board (RX6, Tucker-Davis Techonologies, Alachua, FL, USA) at a sampling rate of 195.3125 kHz. The sound stimuli were attenuated using a programmable attenuator (PA5, Tucker-Davis Techonologies). Click stimuli were generated as a single half-wave of a 5 kHz sine tone. Miniature earphones were inserted into the owl’s ear canals and fixed to a headplate. Acoustic stimuli were fed to these earphones. Stimulus delivery was triggered by National Instruments equipment (NI USB-6259 and BNC-2090A, National Instruments Inc, Austin, TX, USA), and stimulus presentation times were recorded along with the responses. Trigger pulses were configured in MATLAB through Ephus software (Vidrio Technologies LLC, Ashburn, VA, USA). Responses were averaged over 10 repetitions.

### Multi-compartment model of axons

We modeled axons using NEURON (Hines and Carnevale, 1997; Hines et al., 2009) and extended previous work by Kuba and Ohmori (2009), which included the high- and low-threshold potassium channels used by Rathouz and Trussell (1998). The axon was modeled as a sequence of active nodes and passive myelinated segments. The nodes and myelinated segments had lengths of 2 μm and 75 μm, respectively. We included branching axons in our simulations. Branches were generated by connecting two passive segments to a node, and continuing the alternation of active and passive segments in each resulting branch. In cases where the positions of bifurcations or terminations of axons were fixed, the closest passive segment was shortened in order to obtain the total length before the bifurcation or termination.

To evoke an action potential at a designated time, a special conductance was temporarily activated in the first node of Ranvier. The conductance had a reversal potential of 0 mV, a maximal amplitude of 0.05 μS, and a time course described by an alpha-function with time constant 0.01 ms. Soma and axon initial segment were not modeled explicitly.

For the simplified axon geometries used in Figure 1, the branching pattern was fixed as described in the caption, with the exception of the axial position of the branching points in Figure 1E, where a random offset between branching points was drawn from a gamma distribution with mean 400 μm and standard deviation 300 μm. The initial branching point for each axon was offset from the original location by a distance drawn from a Gaussian distribution with mean zero and a width of 300 μm. This was done to smooth out the effects of individual branchings or terminations.

For the axons in Figure 2, branching patterns were generated procedurally, starting with a root segment. In order to avoid artifacts from the stimulus and to simulate a long fiber tract prior to the terminal zone, a sequence of 10 active and passive segments without bifurcations was assumed before the terminal zone. To this root, segments were appended iteratively. Before adding a segment, a decision whether to branch or terminate an axon was drawn from a probability distribution that was dependent on the axial position of the end of the previous segment. These probability distributions were modeled as logistic functions with the parameters adjusted to roughly match the numbers of branchings and terminations shown by Carr and Konishi (1990). Thus, an initial phase dominated by bifurcations was followed by a phase dominated by terminations, with the probability of termination reaching 100% at the end of the terminal zone. The distribution of bifurcations had its maximum at axial location *z* = 0 with a standard deviation of 200 μm. The distribution of terminations had its maximum at at *z* = 500 pm, with a standard deviation of 100 μm. The branching angle had an average of 20°, with a standard deviation of 5°. At branching points, the plane containing the branches had a uniform random orientation, resulting in a 3-dimensional structure of the axon bundle.

Numerical simulations of action potentials propagating along axons yielded the membrane currents from which we calculated extracellular fields. This procedure is described in detail by Gold et al. (2006). Briefly, the extracellular medium is assumed to be a homogeneous volume conductor with conductivity *σ_e_* and a quasi-static approximation of the electrical field potential *ϕ* is made. The extracellular potential *ϕ*(*r*, *t*) at location r and time *t* due to a membrane current density distribution *i*(r,*t*) is then governed by the equation 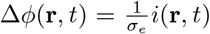, with Δ denoting the Laplace operator (Nunez and Srinivasan, 2006). If the currents *I* are constrained to a volume *W*, this equation has the solution

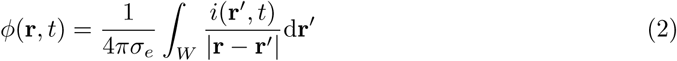

Since the majority of the current flows through the small nodes of Ranvier in myelinated axons, we used the point-source approximation; we did not use the line-source approximation (Holt and Koch, 1999).

Analysis of the resulting extracellular potential included filtering. All filtering was performed with third-order Butterworth filters. The multiunit activity (MUA) was calculated by high-pass filtering the signal with a cutoff of 2500 Hz, setting all samples with negative values to zero, and then low-pass filtering the resulting response with a cutoff of 500 Hz. The LFP was calculated by low-pass filtering the signal with a cutoff of 1000 Hz.

### Mean field model of an axon bundle

To better understand the processes leading to the complex spatio-temporal patterns of extracellular fields, we devised an analytically tractable model of axon bundles. We defined the spatial dimension in cylindrical coordinates r = (*ρ*, *φ*, *z*), and we considered a simple model axon bundle that extended infinitely in the axial *z*-direction at *ρ* = 0. The bundle had a variable number of fibers along the *z* axis, denoted by *n*(*z*), each of which cylindrical with an identical radius ***a.*** This meant that the total cross-sectional area ***A*** of the bundle at a given depth *z* was given by *A*(*z*) = *πa*^2^*n*(*z*). We assumed the axons to be perfect transmission lines, meaning that the action potential is a traveling wave with velocity *v* along the axon. In particular, we neglected delays and distortions that can be induced when an axon branches or terminates. In this case, we could assume that the membrane voltage was the same in each fiber for a given *z* coordinate. From linear cable theory (e.g. Jack et al., 1975), we then obtained the following expression for the total transmembrane current per unit length *I*(*z,t*) from a given membrane potential *V*(*z*,*t*):

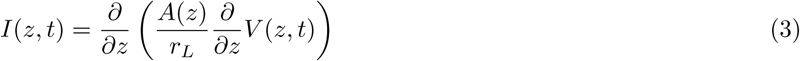

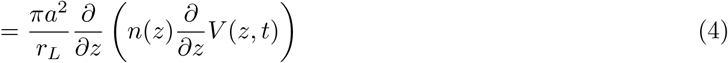

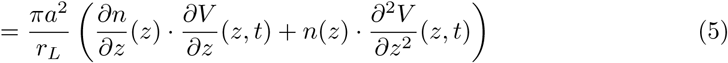

We next calculated the corresponding extracellular field potential *ϕ*(r, *t*) of a given membrane potential waveform *V*(*z*, *t*) propagating through the axon bundle. Due to the rotational symmetry of the system and the fact that current flows only at *ρ* = 0, and the membrane current can be described as 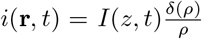, where 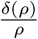 is the Dirac delta distribution for a line at *ρ* = 0.

Applying this membrane current to Equation 2, we obtained

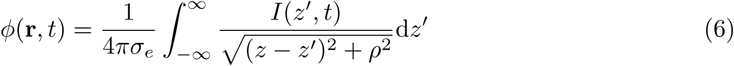

which is independent of the angle *φ*.

To derive the frequency responses in Figure 3, we simulated the membrane potentials as sinusoids, i.e. *V*(*z*,*t*) ***=*** sin (2*πf* · (*z* – *tv*)), with varying frequencies *f* between 100 Hz and 5 kHz and calculated the standard deviation of the response for each frequency individually. The amplitude was the same for all frequencies.

To derive the dipole moment of a simplified projection zone, we considered an axon bundle in which identical spikes with the waveform *V*_spike_(*z*, *t*) propagate as traveling waves with a velocity *v* in positive *z* direction: *V*_spike_(*z*, *t*) ***=*** *V*_spike_(*z* – *tv,* 0). If each of the fibers is stimulated with an inhomogeneous Poisson process, with the driving rate *λ*(*t*) shared among all axons, the average membrane potential across fibers will be *V*(*z,t*) ***=*** *V*_spike_(*z*,*t*) * *λ*(*t*), where * denotes the convolution with respect to the time *t*. Plugging this into Equation 5, we obtained

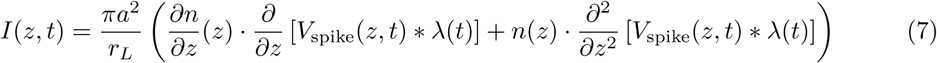

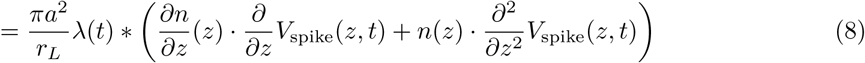

Assuming Gaussian shapes for the firing-rate pulse 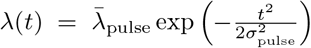, the spike 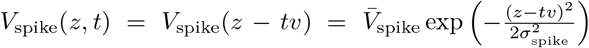, and the fiber bundle projection zone 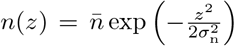, we were able to take advantage of the fact that the product and the convolution of two Gaussians are again Gaussian, and obtained

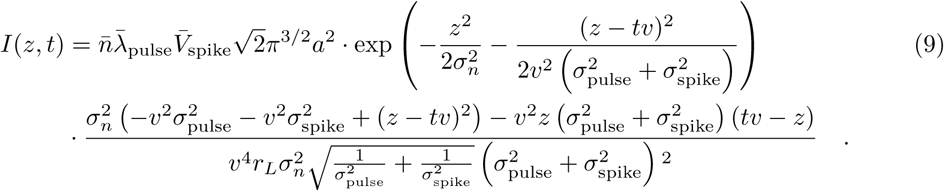

The dipole moment *p*(*t*) is defined as

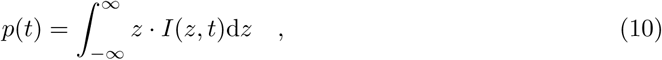

into which we can enter Equation 9 to obtain

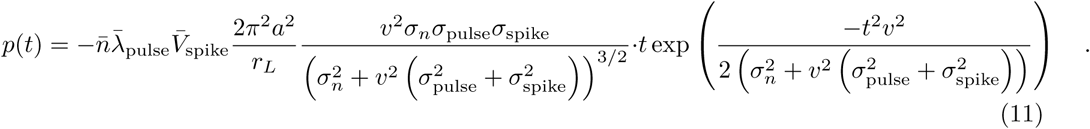

The dipole moment has its maximum amplitude at 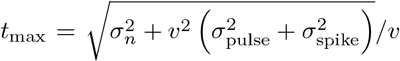, and takes the value

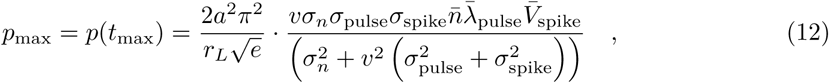

which is presented as Equation 1 in the Results section.

### Model fitting to experimental data

In order to relate the model to experimentally obtained data as shown in Figure 5, we performed a nonlinear least squares fit, minimizing the mean squared error *∊* between the measured potential *ϕ*_measured_ and the model prediction *ϕ*_model_ in Equations 6 and 9 for ***N*** = 32 measurement locations *z_n_* (*n* ***=*** 1,…,*N*) and *M* = 600 time points 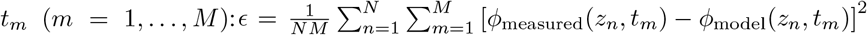. The separation between electrodes was given by the electrode layout as 50 μm. The time between sampling points was 5.12 μs. We achieved the minimization of the error *∊* using the “optimize.minimize” routine provided by the SCIPY package (Jones et al., 2001). The free parameters to be determined by the optimization routine were the distance *ρ*, the velocity *v*, the number of fibers per unit length *n*(*z_n_*) for each measurement location *z_n_*, and the spatial derivative of the average membrane potential 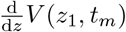 at electrode location *z*_1_ for each time point *t_m_*. We fit the first derivative of the membrane potential in order to better capture the low-frequency components that we found in Figure 1 E, and because the membrane potential only appears as the derivative in the model. The derivative of the membrane voltage at the other locations than *z*_1_ was then determined by the traveling-wave assumption: 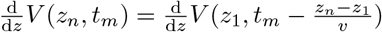, using a linear interpolation between timepoints. The model assumption of a single line of axons with electrodes at a fixed distance is a simplification of a three-dimensional axon tree where the fibers are distributed at various distances. The distance parameter *ρ* in Equation 6 can be interpreted as an average distance in this simplification.

To aid the convergence of the fit algorithm, an initial guess for the number of fibers *n*(*z_Z_*) was set by hand. Initializing the guess to a constant or a fully random number of fibers resulted in a failure to converge. However, Gaussian-like initial guesses converged to a single solution, meaning that the specific initial guess did not alter the final fit result. The results shown in Figure 5 were obtained with an initial guess of a Gaussian with amplitude 12, centered at penetration depth 725 μm with standard deviation 400 μm.

Because of the linearity of Equations 2-6 both in the current *I* and the membrane potential *V*, inferring the membrane voltage *V* from the average over trials of the EFP *ϕ* produces the average membrane voltage *V*. This in turn is the membrane voltage response of a single spike convolved with the peri-stimulus time histogram (PSTH).

## Acknowledgements

This work was supported by the German Federal Ministry for Education and Research Grants 01GQ1001A and 01GQ0972, NIH DCD 000436 and US-American Collaboration in Computational Neuroscience “Field Potentials in the Auditory System” as part of the NSF/NIH/ANR/BMBF/BSF Collaborative Research in Computational Neuroscience Program, 01GQ1505A.

The authors wish to thank Anna Kraemer for help with animal handling and surgery; and Martina Michalikova and Tiziano D’Albis for helpful comments on a draft of this manuscript.

